# Combining RNA-seq data and homology-based gene prediction for plants, animals and fungi

**DOI:** 10.1101/219287

**Authors:** Jens Keilwagen, Frank Hartung, Michael Paulini, Sven O. Twardziok, Jan Grau

## Abstract

**Motivation:** Genome annotation is of key importance in many research questions. The identification of protein-coding genes is often based on transcriptome sequencing data, ab-initio or homology-based prediction. Recently, it was demonstrated that intron position conservation improves homology-based gene prediction, and that experimental data improves ab-initio gene prediction.

**Results:** Here, we present an extension of the gene prediction tool GeMoMa that utilizes amino acid sequence conservation, intron position conservation and optionally RNA-seq data for homology-based gene prediction. We show on published benchmark data for plants, animals and fungi that GeMoMa performs better than the gene prediction programs BRAKER1, MAKER2, and CodingQuarry, and purely RNA-seq-based pipelines for transcript identification. In addition, we demonstrate that using multiple reference organisms may help to further improve the performance of GeMoMa. Finally, we apply GeMoMa to four nematode species and to the recently published barley reference genome indicating that current annotations of protein-coding genes may be refined using GeMoMa predictions.

**Availability:** GeMoMa has been published under GNU GPL3 and is freely available at http://www.jstacs.de/index.php/GeMoMa.

## 1 Introduction

The annotation of protein-coding genes is of critical importance for many fields of biological research including, for instance, comparative genomics, functional proteomics, gene targeting, genome editing, phylogenetics, transcriptomics, and phylostratigraphy. The process of annotating protein-coding genes to an existing genome (assembly) can be described as specifying the exact genomic location of genes comprising all (partially) coding exons. A difficulty in gene annotation is distinction between protein-coding genes, transposons and pseudogenes.

Genome annotation pipelines utilize three main sources of information, namely evidence from wet-lab transcriptome studies (Trapnell *et al.*, 2010; Pertea *et al.*, 2015), ab-initio gene prediction based on general features of (protein-coding) genes (Solovyev *et al.*, 2006; Stanke *et al.*, 2008), and homology-based gene prediction relying on gene models of (closely) related, well-annotated species (Slater and Birney, 2005; She *et al.*, 2011; Keilwagen *et al.*, 2016).

Experimental data allow for inferring coverage of gene predictions and splice sites bordering their exons, which may assist computational ab-initio or homology-based approaches. Due to the progress in the field of next generation sequencing, RNA-seq has revolutionized transcriptomics (Wang *et al.*, 2009). Today, RNA-seq data is available for a wide range of organisms, tissues and environmental conditions, and can be utilized for genome annotation pipelines.

In recent years, several programs have been developed that combine multiple sources allowing for a more accurate prediction of proteincoding genes (Holt and Yandell, 2011; Testa *et al.*, 2015; Hoff *et al.*, 2016). MAKER2 is a pipeline that integrates support of different resources including ab-initio gene predictors and RNA-seq data (Holt and Yandell, 2011). CodingQuarry is a pipeline for RNA-Seq assembly-supported training and gene prediction, which is only recommended for application to fungi (Testa *et al.*, 2015). Recently, Hoff et al. (2016) published BRAKER1 a pipeline for unsupervised RNA-seq-based genome annotation that combines the advantages of GeneMark-ET (Lomsadze *et al.*, 2014) and AUGUSTUS (Stanke *et al.*, 2008).

Here, we present an extension of GeMoMa (Keilwagen *et al.*, 2016) that utilizes RNA-seq data in addition to amino acid sequence and intron position conservation. We investigate the performance of GeMoMa on publicly available benchmark data (Hoff *et al.*, 2016) and compare it with state-of-the-art competitors (Holt and Yandell, 2011; Testa *et al.*, 2015; Hoff *et al.*, 2016).

Subsequently, we demonstrate how combining homology-based predictions based on gene models from multiple reference organisms can be used to improve the performance of GeMoMa. Finally, we apply GeMoMa to four nematode species provided by Wormbase (Howe *et al.*, 2016) and to the recently published barley reference genome (Mascher *et al.*, 2017), where GeMoMa predictions will be included into future versions of the corresponding genome annotations.

## 2 Methods

In this section, we describe recent extensions of GeMoMa to make use of evidence from RNA-seq data, the RNA-seq pipelines used and the data considered in the benchmark and application studies.

### 2.1 GeMoMa using RNA-seq

GeMoMa predicts protein-coding genes utilizing the general conservation of protein-coding genes on the level of their amino acid sequence and on the level of their intron positions, i.e., the locations of exon-exon boundaries in CDSs (Keilwagen *et al.*, 2016). To this end, sequences of (partially) protein-coding exons are extracted from well-annotated reference genomes. Individual exons are then matched to loci on the target genome using tblastn (Altschul *et al.*, 1990), matches are adjusted for proper splice sites, start codons and stop codons, respectively, and joined to full, protein-coding genes models. In this process, the conserved dinucleotides GT and GC for donor splice sites, and AG for acceptor splice sites have been used for the identification of splice sites bordering matches to the (partially) protein-coding exons of the reference transcripts. The improved version of GeMoMa may now also include experimental splice site evidence extracted from mapped RNA-seq data to improve the accuracy of splice site and, hence, exon annotation. We visualize the extended GeMoMa pipeline in Fig. S1.

Starting from mapped RNA-seq data, the module *Extract RNA-seq evidence* (ERE) allows for extracting introns and, if user-specified, read coverage of genomic regions. GeMoMa filters these introns using a user-specified minimal number of split reads within the mapped RNA-seq data. Introns passing this filter define donor and acceptor splice sites, which are treated independently within GeMoMa. If splice sites with experimental evidence have been detected in a genomic region with a good match to an exon of a reference transcript, these are collected for further use. If no splice sites with experimental evidence have been detected in a genomic region with a good match to an exon of a reference transcript, GeMoMa resorts to conserved dinucleotides allowing to identify gene models that are not covered by RNA-seq data due to, e.g., very specifically or lowly expressed transcripts. Combining two potential exons, all in-frame combinations using the collected donor and acceptor splice sites are tested and scored according to the reference transcript. The best combination is used for the prediction.

Based on this experimental evidence, the improved version of GeMoMa provides several new properties reported for gene predictions. The most prominent features are *transcript intron evidence* (tie) and *transcript percentage coverage* (tpc). The tie of a transcript varies between 0 and 1, and corresponds to the fraction of introns (i.e., splice sites of two neighboring exons) that are supported by split reads in the mapped RNA-seq data. In case of transcripts comprising a single coding exon, NA is reported. The tpc of a transcript also varies between 0 and 1, and corresponds to the fraction of (coding) bases of a predicted transcript that are also covered by mapped reads in the RNA-seq data.

GeMoMa allows for computing and ranking multiple predictions per reference transcript, but does not filter these predictions. Predictions of different reference transcripts might be highly overlapping or even identical, especially if the reference transcripts are from the same gene family. Since GeMoMa 1.4, the default parameters for number of predictions and contig threshold have been changed which might lead to an increased number of highly overlapping or identical predictions. In addition, it might be beneficial to run GeMoMa starting from multiple reference species to broaden the scope of transcripts covered by the predictions. However, these may also result in redundant predictions for, e.g., orthologs or paralogs stemming from the different reference species considered. To handle such situations, the new module *GeMoMa annotation filter* (GAF) of the improved version of GeMoMa now allows for joining and reducing such predictions using various filters. Filtering criteria comprise the relative GeMoMa score of a predicted transcript, filtering for complete predictions (starting with start codon and ending with stop codon), and filtering for evidence from multiple reference organisms. In addition, GAF also joins duplicate predictions that originate from different reference transcripts.

Initially, GAF filters predictions based on their relative GeMoMa score, i.e., the GeMoMa score divided by the length of the predicted protein. This filter removes spurious predictions. Subsequently, the predictions are clustered based on their genomic location. Overlapping predictions on the same strand yield a common cluster. For each cluster, the prediction with the highest GeMoMa score is selected. Non-identical predictions overlapping the high-scoring prediction with at least a user-specified percentage of borders (i.e., splice sites, start and stop codon, cf. *common border filter*) are treated as alternative transcripts. Predictions that have completely identical borders to any previously selected prediction are removed and only listed in the GFF attribute field *alternative*. All filtered predictions of a cluster are assigned to one gene with a generic gene name. Finally, GAF checks for nested genes in the cluster looking for discarded predictions that do not overlap with any selected prediction, which are recovered.

In addition to the modules for annotating a genome (assembly) described above, we also provide two additional modules in GeMoMa for analyzing and comparing to prediction to a reference annotation. The module *CompareTranscripts* determines that CDS of the reference annotation with the largest overlap with the prediction utilizing the *F*_1_ measure as objective function (Keilwagen *et al.*, 2016). The module *AnnotationEvidence* computes tie and tpc of all CDSs of a given annotation. Hence, these two modules can be used to determine, whether a prediction is known, partially known or new and whether the overlapping annotation has good RNA-seq support.

### 2.2 MAKER2 predictions

Recently, we have shown that GeMoMa outperforms state-of-the-art homology-based gene predictors (Keilwagen *et al.*, 2016). We are not aware of any homology-based gene prediction program that allows for incorporating of RNA-seq data. Hence, we provide predictions of MAKER2 using the same reference proteins as GeMoMa for a minimal comparison. Internally, MAKER2 uses exonerate (Slater and Birney, 2005) for homology-based gene prediction. We run MAKER2 with default parameters except `protein2genome=1`, and `genome` and protein set to the respective input files. In addition, we run MAKER2 using (i) RNA-seq data in form of Trinity 2.4 transcripts (-jaccard clip) (Haas *et al.*, 2013), (ii) homology in form of proteins of one related reference species, and (iii) ab-initio gene prediction in form of Augustus 3.3 (Stanke *et al.*, 2008). In this case, we run MAKER2 with default parameters except `genome, est, protein`, and `augustus_species`, which have been set to the corresponding species. For comparison, we run Maker2 with the same parameter settings but using the GeMoMa predictions for `protein_gff` instead of using `protein`.

### 2.3 RNA-seq pipelines

Computational pipelines have been used to infer gene annotation from RNA-seq data produced by next generation sequencing methods. Dozens of tools and tool combinations have been proposed. Here, we focus on the short read mapper TopHat2 (Kim *et al.*, 2013), the transcript assemblers Cufflinks (Trapnell *et al.*, 2010) and StringTie (Pertea *et al.*, 2015), and the coding sequence predictor TransDecoder (Haas *et al.*, 2013). Based on the transcript assemblers, we build two RNA-seq pipelines following the instructions in Hoff et al. (2016).

### 2.4 Data

For the benchmark studies, we consider target species and their genome versions as specified in the BRAKER1 supplement. For the homology-based prediction by GeMoMa, we choose one closely related reference species per target species that are sequenced and annotated (Rawat *et al.*, 2015; Howe *et al.*, 2016; Matthews *et al.*, 2015; Rhind *et al.*, 2011). For these species, we consider the latest genome versions available (Tab. S1). For the analysis of *C. elegans*, we use the manually curated gene set of *C. briggsae* provided by Wormbase. In addition, we use the experimental evidence from RNA-seq data referenced in the BRAKER1 publication.

For the analysis of the four nematode species, *C. brenneri*, *C. briggsae*, *C. japonica*, and *C. remanei*, we use the genome assembly and gene annotation of Wormbase WS257 (Howe *et al.*, 2016). We choose the model organism *C. elegans* as reference species (Tab. S2). In addition to genome assembly and gene annotation, we also use publicly available RNA-seq data of these four nematode species, which have been mapped by Wormbase using STAR (Dobin *et al.*, 2013). We used a minimum intron size of 25 bp, a maximum intron size of 15Kb, specify that only reads mapping once or twice on the genome are reported, and alignments are reported only if their ratio of mismatches to mapped length is less than 0.02. In accordance with the previous benchmark study, we use the manually curated gene set of Wormbase.

For the analysis of barley, we use the latest genome assembly and gene annotation (Mascher *et al.*, 2017). As reference species, we choose *A. thaliana* (Lamesch *et al.*, 2012), *B. distachyon* (International Brachypodium Initiative, 2010), *O. sativa* (Ouyang *et al.*, 2007), and *S. italica* (Bennetzen *et al.*, 2012) (Tab. S3). In addition to genome assembly and gene annotation, we also used RNA-seq data from four different public available data sets (ERP015182, ERP015986, SRP063318, SRP071745). Reads were mapped and assembled using Hisat2 and StringTie (Pertea *et al.*, 2016). As reference annotation, we used the union of high and low confidence annotation.

As independent evidence for validating GeMoMa predictions in the nematode species and barley, we use ESTs and cDNAs. While Wormbase provides coordinates for *best BLAT matches*, we adapt the pipeline and download all available EST from NCBI and map them to the genome using BLAT (Kent, 2002).

## 3 Results and Discussion

### 3.1 Benchmark

The comparison of different software pipelines is often critical as a) specific parameters settings might be crucial for good results and b) different input might be used. For these reasons, we designed the benchmark as follows. First, we use publicly available gene predictions results. Second, we limit the number of reference species to one in the initial study.

We used GeMoMa for predicting the gene annotations of *A. thaliana*, *C. elegans*, *D. melanogaster*, and *S. pombe*. In Table 1, we summarize the performance of BRAKER1, MAKER2, and CodingQuarry as reported in Hoff et al. (2016), as well as the performance of GeMoMa with and without RNA-seq evidence, purely RNA-seq-based pipelines and various MAKER2 predictions. For all comparisons, we provide sensitivity (Sn) and specificity (Sp) for the categories gene, transcript, and exon, respectively (Keibler and Brent, 2003).

**Table 1:** Benchmark results on the BRAKER1 test sets. The target species are given in multi-column rows. The same reference species, which is given in brackets, is used for all tools using homology-based gene prediction indicated by plus. The asterisks indicates that the performance of BRAKER1, MAKER2 and CodingQuarry is given as reported in Hoff et al. (2016). The highest value per line is depicted in bold-face.

|  | MAKER2<sup>+</sup> (exonerate) | GeMoMa<sup>+</sup> without RNA-seq data | GeMoMa<sup>+</sup> with RNA-seq data | RNAseq-Cufflinks | RNAseq-StringTie | BRAKER1* | MAKER2* | CodingQuarry* | MAKER2<sup>+</sup> (exonerate, Trinity, Augustus) | MAKER2<sup>+</sup> (GeMoMa, Trinity, Augustus) |
| --- | --- | --- | --- | --- | --- | --- | --- | --- | --- | --- |
| *Arabidopsis thaliana* (ref. *A. lyrata*) |  |  |  |  |  |  |  |  |  |  |
| Gene Sn | 44.0 | 61.3 | **66.5** | 28.9 | 35.9 | 64.4 | 51.3 | NA | 56.9 | 57.9 |
| Gene Sp | 47.8 | 65.7 | **71.3** | 47.9 | 59.1 | 52.0 | 52.5 | NA | 65.7 | 67.8 |
| Transcript Sn | 37.5 | 52.2 | **57.2** | 26.6 | 33.7 | 55.0 | 43.5 | NA | 48.3 | 49.1 |
| Transcript Sp | 47.8 | 65.7 | 65.3 | 35.6 | 48.3 | 50.9 | 52.5 | NA | 65.7 | **67.8** |
| Exon Sn | 70.0 | 79.3 | 80.6 | 58.1 | 60.8 | 82.9 | 76.1 | NA | 81.8 | **82.1** |
| Exon Sp | 81.9 | 86.6 | 87.5 | 81.9 | 87.1 | 79.0 | 76.1 | NA | 87.5 | **88.6** |
| *Caenorhabditis elegans* (ref. *C. briggsae*) |  |  |  |  |  |  |  |  |  |  |
| Gene Sn | 26.2 | 39.6 | 49.1 | 18.7 | 22.6 | **55.0** | 41.0 | NA | 40.5 | 47.3 |
| Gene Sp | 38.0 | 49.9 | **63.8** | 29.1 | 36.1 | 55.2 | 30.8 | NA | 51.5 | 56.4 |
| Transcript Sn | 21.0 | 30.7 | 39.8 | 16.2 | 20.0 | **43.0** | 31.3 | NA | 31.4 | 36.2 |
| Transcript Sp | 38.0 | 49.9 | **58.7** | 24.1 | 30.1 | 53.2 | 30.8 | NA | 51.5 | 56.4 |
| Exon Sn | 50.3 | 64.2 | 67.1 | 54.4 | 59.1 | **80.2** | 69.4 | NA | 70.5 | 75.2 |
| Exon Sp | 82.6 | 81.5 | **87.5** | 81.3 | 84.1 | 85.3 | 62.3 | NA | 85.6 | 86.7 |
| *Drosophila melanogaster* (ref. *D. simulans*) |  |  |  |  |  |  |  |  |  |  |
| Gene Sn | 64.3 | 78.2 | **83.1** | 55.7 | 55.2 | 64.9 | 55.2 | NA | 61.5 | 64.0 |
| Gene Sp | 69.2 | 81.6 | **87.1** | 71.3 | 73.5 | 59.4 | 46.3 | NA | 69.6 | 71.9 |
| Transcript Sn | 44.1 | 52.9 | **65.0** | 48.7 | 49.0 | 46.1 | 38.5 | NA | 42.7 | 44.3 |
| Transcript Sp | 69.2 | **81.6** | 81.2 | 60.1 | 65.7 | 57.9 | 46.3 | NA | 69.6 | 71.9 |
| Exon Sn | 69.0 | 76.3 | **80.0** | 67.8 | 66.2 | 75.0 | 66.5 | NA | 74.3 | 76.3 |
| Exon Sp | 89.1 | 92.0 | **93.3** | 85.4 | 88.3 | 81.7 | 66.9 | NA | 88.0 | 89.1 |
| *Schizosaccharomyces pombe* (ref. *S. octosporus*) |  |  |  |  |  |  |  |  |  |  |
| Gene Sn | 49.2 | 76.4 | 79.2 | 69.0 | 65.8 | 77.4 | 42.8 | **79.7** | 71.6 | 74.6 |
| Gene Sp | 59.9 | 84.6 | 88.0 | 93.8 | 92.5 | 80.5 | 68.7 | 72.6 | 88.1 | **89.1** |
| Transcript Sn | 49.2 | 76.4 | 79.2 | 69.0 | 65.8 | 77.4 | 42.8 | **79.7** | 71.6 | 74.6 |
| Transcript Sp | 59.9 | 84.6 | 87.6 | 80.5 | 71.3 | 76.5 | 68.7 | 72.6 | 88.1 | **89.1** |
| Exon Sn | 56.1 | 81.6 | 83.1 | 77.2 | 77.7 | **83.2** | 50.1 | 79.6 | 79.2 | 81.2 |
| Exon Sp | 73.3 | 88.6 | 91.9 | 87.6 | 81.7 | 83.2 | 71.4 | 81.7 | 92.0 | **92.6** |

First, we compare the two purely homology-based predictions, namely on the one hand side MAKER2 using exonerate and on the other hand side GeMoMa without RNA-seq data. In all cases, we use the same reference species and reference proteins. We find that MAKER2 using only homologous proteins has a higher exon specificity than GeMoMa without RNA-seq data for *C. elegans*, while the opposite is true for all other categories and target species.

Second, we additionally consider RNA-seq data. MAKER2 does not allow for combining RNA-seq evidence and homology-based predictions without using any ab-initio gene predictor. In contrast, GeMoMa allows for additionally using intron position conservation and RNA-seq data. For this reason, we compare the performance of GeMoMa with and without RNA-seq evidence (Table 1). We find that sensitivity and specificity in almost all cases increases by up to 13.9 with only two exceptions for transcript specificity of *A. thaliana* and *D. melanogaster* which decreases by at most 0.4. Hence, we summarize that RNA-seq evidence improves the sensitivity and specificity of GeMoMa and should be used if available.

Third, we compare the performance of GeMoMa using RNA-seq evidence to that of purely RNA-seq-based pipelines, namely Cufflinks and StringTie (Table 1). We find for all four species that GeMoMa using RNA-seq evidence outperforms purely RNA-seq-based pipelines. Interestingly, purely RNA-seq-based pipelines also yield the worst gene/transcript sensitivity and specificity for *C. elegans*.Comparing the results based on different transcript assemblers, we find that the results based on StringTie are better than those based on Cufflinks for *A. thaliana* and *C. elegans*, while the opposite is true for *S. pombe*. For *D. melanogaster*, both pipelines perform comparably. Additional RNA-seq reads increasing the coverage might improve the performance of purely RNA-seq-based pipelines but could also improve the performance of GeMoMa.

Summarizing these three observations, we find that GeMoMa performs better than purely homology-based or purely RNA-seq-based pipelines and that including RNA-seq data improves the performance of GeMoMa.

Hence, we compare GeMoMa to combined gene prediction approaches. Specifically, we compare the performance of GeMoMa using RNA-seq evidence to BRAKER1 in Fig. 1, which provides the best overall performance in Hoff et al. (2016). We find that GeMoMa performs better than BRAKER1 for the categories gene and transcript with the exception of gene and transcript sensitivity for *C. elegans*. Interestingly, we find the biggest improvements for *D. melanogaster* where gene/transcript sensitivity and specificity increases between 18.2 and 27.7. For the exon category, we find a less clear picture. In total, we observe the worst results for *C. elegans* where the sensitivity for all three categories decreases between 3.2 and 13.2, while the specificity increases only between 2.2 and 8.6. Notably, we generally find the worst gene/transcript sensitivity and specificity for *C. elegans* compared with the other target species considering the best performance of all tools.

**Figure 1:**
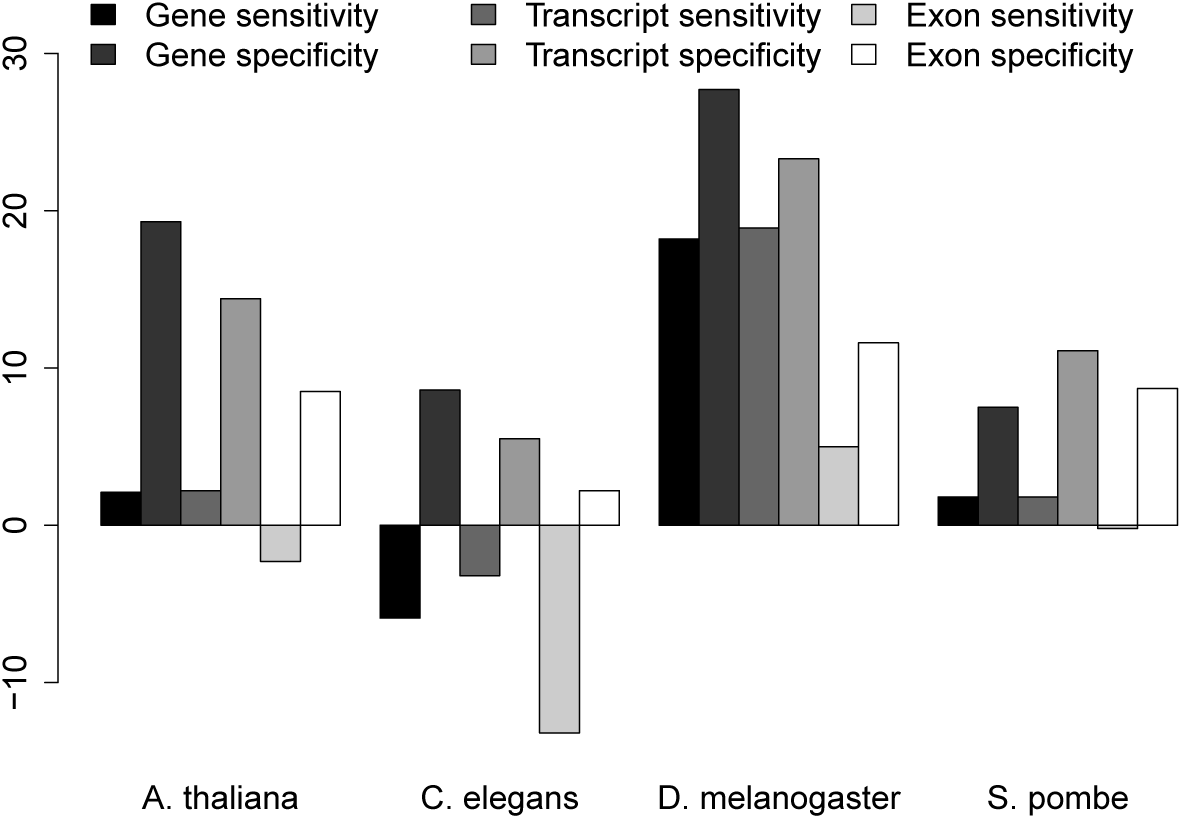
Benchmark results. The y-axis depicts the difference between the GeMoMa with RNA-seq data and the BRAKER1 performance.

In summary, we find that the gene predictors MAKER2, BRAKER1, CodingQuarry and GeMoMa, and the transcript assemblers Cufflinks and StringTie often perform quite well on exon level. The main difference becomes evident on transcript and gene level, where exons need to be combined correctly (Table 1) as reported earlier (Steijger *et al.*, 2013; Conesa *et al.*, 2016). Homology-based gene predictors might benefit from experimentally validated and manually curated reference transcripts guiding the prediction of transcripts in the target organism.

Although GeMoMa performed well, it is not able to predict genes that do not show any homology to a protein in the reference species, while ab-initio gene predictors might fail in other cases. As both types of approaches have their specific advantages, users will probably use combinations of different gene predictors in practice to obtain a comprehensive gene annotation.

### 3.2 Combined gene prediction pipelines

Combined gene prediction pipelines, as for instance MAKER2, use RNA-seq evidence, homology-based and ab-initio methods for predicting final gene models. MAKER2 uses exonerate by default for homology-based gene prediction. However, MAKER2 also provides the possibility to use other homology-based gene predictors instead of exonerate (cf. parameter protein gff). For this reason, we compare the performance of MAKER2 using either exonerate or GeMoMa for homology based gene prediction (Table 1). In addition, we use Augustus as ab-initio gene prediction program and Trinity transcripts in MAKER2. We find that MAKER2 using GeMoMa performs better than MAKER2 using exonerate for all species and all measure. The improvement varies between 0.3% and 6.8% with clearly the biggest improvement for *C. elegans*.

In addition, we find that the MAKER2 performance is substantially improved compared to the performance of the the previously reported MAKER2 predictions, either purely based on proteins (cf. Table 1, column MAKER2^+^ (exonerate)) or as reported in Hoff et al. (2016) (cf. Maker2^∗^). These other predictions do not utilize all available sources of information as they either ignore RNA-seq data and ab-initio gene prediction or homology to proteins of related species. Based on this observation, we agree that combined gene prediction pipelines benefit from the inclusion of all available evidence and that performance is decreased if some important evidence is missed (Holt and Yandell, 2011).

Furthermore, we compare GeMoMa using RNA-seq evidence with MAKER2 using RNA-seq evidence, homology-based and ab-initio gene prediction. In some cases, it is hard to compare these results as sensitivity of one tool is higher than the sensitivity of the other tool and the opposite is true for specificity. In machine learning, recall, also known as sensitivity, and precision, which is called specificity in the context of gene prediction evaluation (Burset and Guigó, 1996), are combined into a single scalar value called F1 measure (Powers, 2011) that can be compared more easily. We combined sensitivity and specificity resulting in an F1 measure for each evaluation level gene, transcript and exon (Table S4) We find that in many cases GeMoMa using RNA-seq evidence outperforms MAKER2. The reason for this observation might be that RNA-seq data and homology based gene prediction is used in MAKER2 to train ab-initio gene predictors, in this case Augustus. With the recommended parameter setting, homology-based gene predictions are not directly used for the final prediction and doing so might further improve performance.

### 3.3 Influence of reference species

Utilizing different fly species from FlyBase (Gramates *et al.*, 2017), we scrutinize the influence of different or multiple reference species on the performance of GeMoMa using RNA-seq data (Tab. S5). In Fig. 2, we depict gene sensitivity and gene specificity for eight different reference species indicated by points. We find that performance varies with the reference species. In this specific case, *D. sechellia* and *D. persimilis* yield the worst results for single reference-based predictions. This observation might be related to the fact that genome assembly of *D. sechellia* and *D. persimilis* is of lower quality (Clark *et al.*, 2007), while the genome of *D. simulans* has been updated (Hu *et al.*, 2013) later. Besides these two outliers, the performance of the different fly species as reference species for *D. melanogaster* in GeMoMa correlates with their evolutionary distance (Singh *et al.*, 2009). Generally speaking, the closer a reference species is related to the target species *D. melanogaster*, the better is the performance in terms of gene sensitivity and specificity. Hence, we speculate that two requirements must be met to have a good reference species. First, the evolutionary distance between reference and target species should be small and second, the genome assembly and annotation of the reference species should be comprehensive and of high quality.

**Figure 2:**
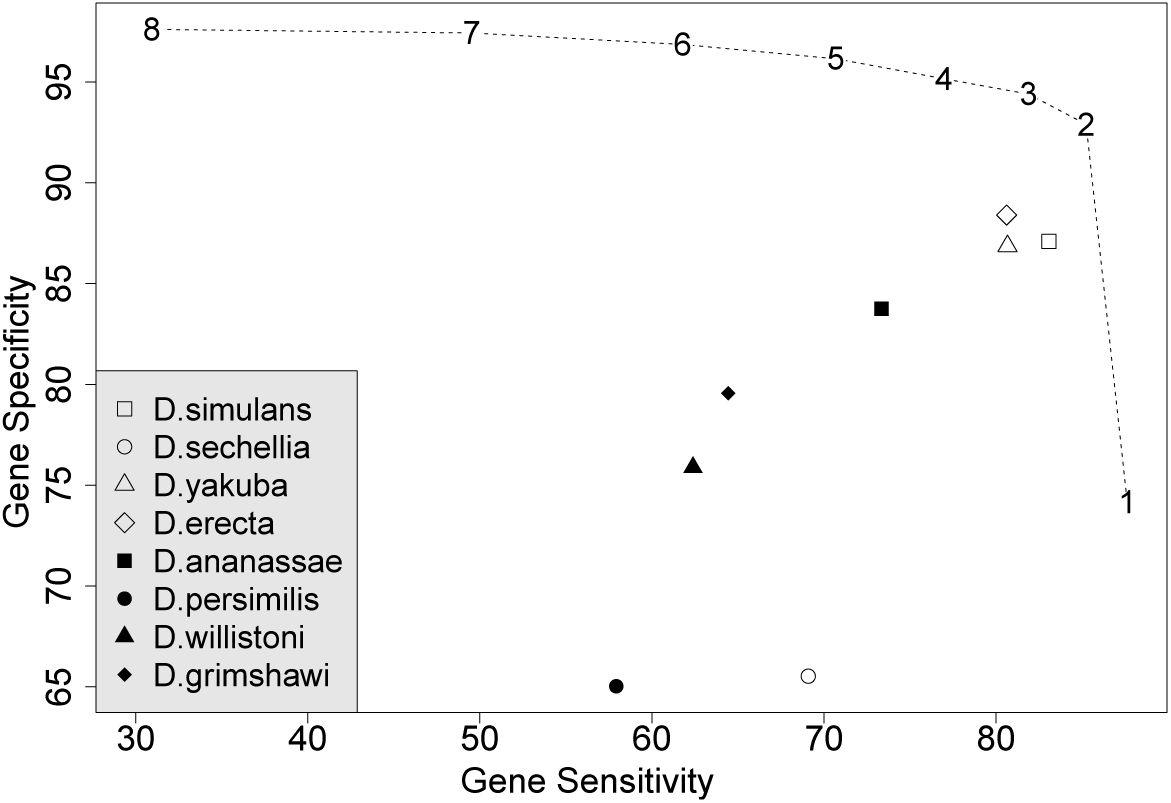
Gene sensitivity and specificity for *D. melanogaster* using different or multiple reference species in GeMoMa. The points correspond to the eight reference species. In addition, the dashed line indicates the usage of multiple reference species. Using multiple reference species allows for filtering identical predictions from several reference as indicated by the numbers.

The new GAF module of GeMoMa allows for combining the predictions based on different reference organisms. The combined predictions may be filtered by number of reference species with perfect support (#evidence), as indicated by the dashed line. We find that combining multiple reference organisms improves prediction performance and stability. Depending on the number of supporting reference organisms required, gene specificity and gene sensitivity may be balanced according to the needs of a specific application. We observe that (i) gene sensitivity increases but specificity decreases when requiring support from at least one reference organism, whereas (ii) gene specificity increases but sensitivity decreases severely filtering for perfect support from all eight reference species. In summary, the inclusion of multiple reference species may yield an improved prediction performance for GeMoMa using the GAF module, where we suggest to filter predictions for support by at least two but not necessarily all reference species.

Furthermore, we check whether GeMoMa allows for identifying new transcripts in *D. melanogaster* that do not overlap with any annotated transcript but are supported by RNA-seq data. First, we check whether we could identify transcripts based on the GeMoMa predictions using *D. simulans* as reference organism. We find 35 multi-coding-exon predictions that do not overlap with any annotated transcript but have a tie of 1, i.e., all introns are supported by split reads in the RNA-seq data (see Methods). In addition, we find 15 single-coding-exon predictions that do not overlap with any annotated transcript but have a tpc of 1, i.e., that are fully covered by mapped RNA-seq reads. Second, we check whether we could identify transcripts that are supported by at least two of the eight reference species (cf. above). We find 14 multi-coding-exon predictions that do not overlap with any annotated transcript, obtain a tie of 1 and are supported by at least two of the eight reference species. In addition, we find 9 single-coding-exon predictions that do not overlap with any annotated transcript, have a tpc of 1 and are supported by at least two of the five reference species. In summary, those genes supported by multiple reference organisms or additional RNA-seq data might be promising candidates for extending the existing genome annotation of *D. melanogaster*.

### 3.4 Analysis of nematode species

The relatively poor results for *C. elegans* in the benchmark study, might be due to insufficiencies in the current *C. briggsae* annotation. Hence, we decided to scrutinize the Wormbase annotation of four nematode species comprising *C. brenneri*, *C. briggsae*, *C. japonica*, and *C. remanei* based on the model organism *C. elegans*. We compare GeMoMa predictions with manually curated CDS from Wormbase. Based on RNA-seq evidence, we collect multi-coding-exon predictions of GeMoMa with tie=1 and compare these to the annotation as depicted in Fig. 3.

**Figure 3:**
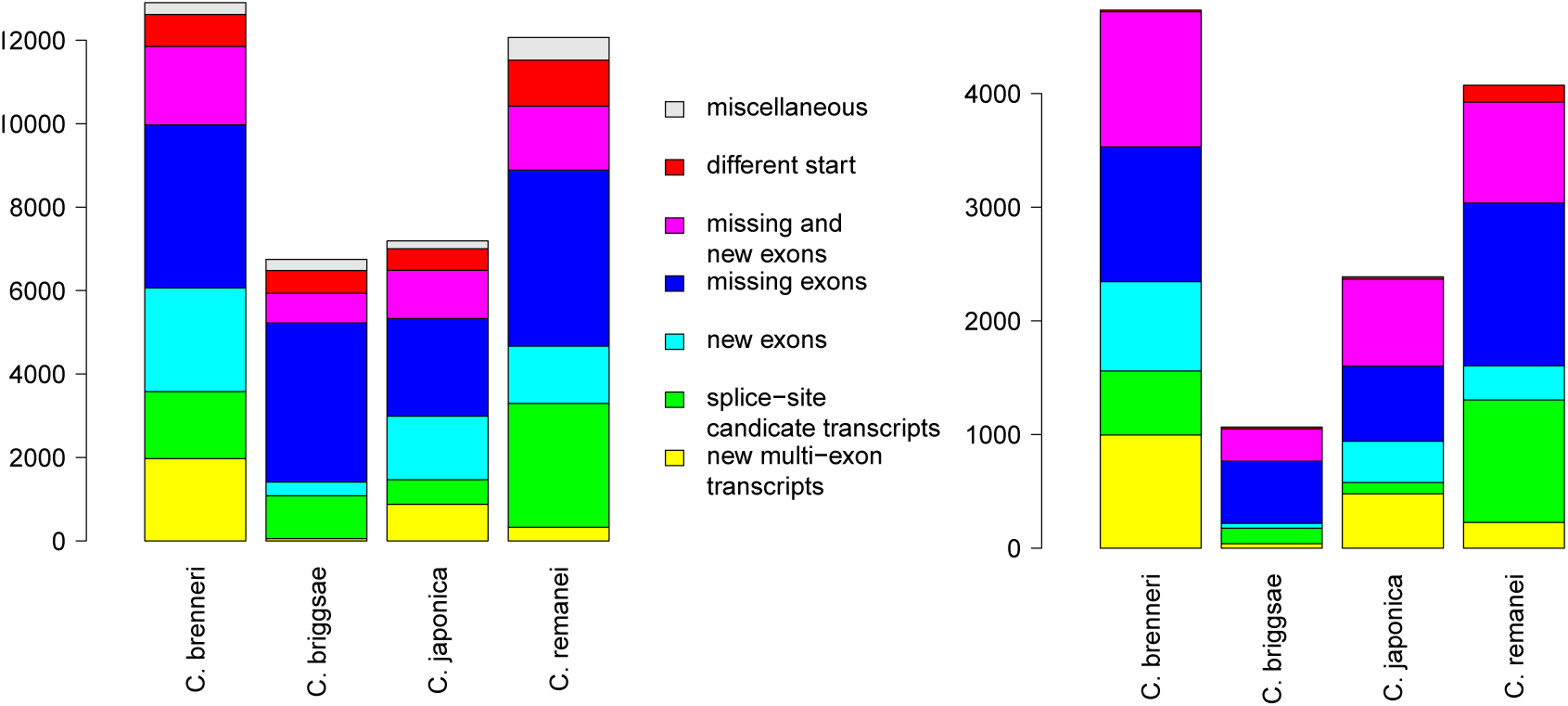
Summary of difference for GeMoMa predictions with tie=1. The relaxed evaluation (left panel) depicts differences between GeMoMa predictions and annotation without any filter on the annotation, while the conservative evaluation (right panel) applies additional filters for the annotation (cf. main text). Predictions that do not overlap with any annotated CDS are depicted in yellow, Predictions that differ from annotated CDSs only in splice sites are depicted in green, predictions that have additional exons compared to annotated CDSs are depicted in turquoise, predictions that missed some exons compared to annotated CDSs are depicted in blue, predictions with additional and missing exons compared to annotated CDSs are depicted in pink, predictions that only differ in the start of the CDS compared to annotated CDS are depicted in red, and any other category is depicted in gray.

In summary, we find between 6 749 differences for *C. briggsae* and 12 903 for *C. brenneri* (cf. Fig. 3(a)). The most interesting category are new multicoding-exon predictions, which vary between 53 for *C. briggsae* and 1974 for *C. brenneri*. The largest category are GeMoMa predictions that missed exons compared to annotated CDSs, which vary between 2340 for *C. japonica* and 4 220 for *C. remanei*.

We additionally filter the transcripts showing differences to obtain a smaller, more conservative set of high-confidence predictions. First, we filter new multi-coding exon GeMoMa predictions for tpc=1 obtaining between 39 and 996 for *C. briggsae* and *C. brenneri*, respectively. Second, we filter GeMoMa predictions that have different splice sites compared to highly overlapping annotated transcripts, contain new exons, have missing exons, or have new and missing exons for tie<1 of the overlapping annotation. We obtain between 100 and 1 079 predictions with different splice-site, between 42 and 786 predictions containing new exons, between 548 and 1 431 predictions with missing exons, and between 284 and 1 191 predictions with new and missing exons. Finally, for GeMoMa predictions that differ in the start codon compared to the annotation, we filter for tpc=1 of the GeMoMa prediction and tpc<1 for the annotation obtaining between 14 and 149 for *C. brenneri* and *C. remanei*, respectively. In summary, we obtain between 1 065 predictions differing from the annotation for *C. briggsae* and 4735 predictions for *C. brenneri*, respectively (cf. Fig. 3(b)) using these strict criteria. Despite the overall reduction of transcripts considered, GeMoMa predictions that missed exons compared to annotated CDSs are the largest category for all four nematode species.

For both evaluations, we find that the predictions for *C. briggsae* are in better accordance with the annotation than the predictions of the remaining three nematode species. One possible explanation might be that the annotation of *C. briggsae* has recently been updated using RNA-seq data (Gary Williams, personal communication), while the annotation of *C. japonica* is based on Augustus (Erich Schwartz, personal communication) and the annotation of the other two nematodes are NGASP sets from multiple ab-initio gene prediction programs (Coghlan *et al.*, 2008). For *C. japonica*, we find the second best results, although *C. japonica* is phylogenetically more distantly related to *C. elegans* than the remaining two nematodes (Kiontke *et al.*, 2011). This is additional evidence that the annotation pipeline employed has a decisive influence on the quality and completeness of the annotation.

In addition, we checked for *C. brenneri* whether the GeMoMa predictions partially overlap with cDNAs or ESTs mapped to the *C. brenneri* genome. In 472 cases, the prediction overlaps with a cDNA or EST, but not with the annotation. In 364 out of these 472 cases, the prediction has tie=1. To evaluate the predictions, we manually checked about 9% (43) of the predicted missing genes with tie=1. Based on RNA-seq data, protein homology, cDNA/ESTs and manual curation, 95% were genuine new isoforms which have been missed in the original *C. brenneri* gene set. This shows that GeMoMa is valuable in finding isoforms missed by traditional prediction methods.

**Table 2:**
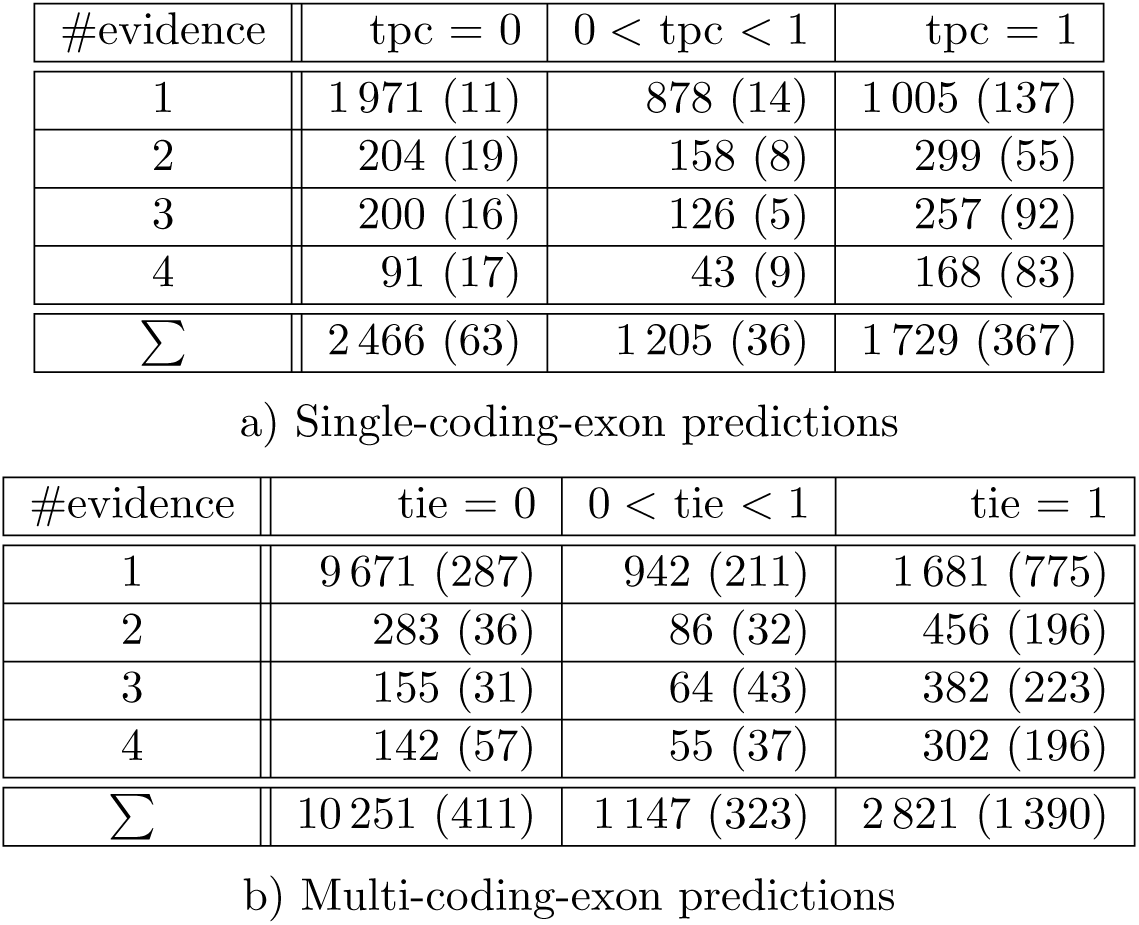
Predictions that do not overlap with any high or low confidence annotation. The numbers in parenthesis depict those predictions that are partially supported by any best BLAT hit of ESTs.

### 3.5 Analysis of barley

Complementary to the studies in animals in the last subsection, we used GeMoMa to predict the annotation of protein-coding genes in barley (*Hordeum vulgare*). Based on the benchmark results for *D. melanogaster*, we used several reference organisms to predict the gene annotation using GeMoMa and GAF and finally obtain 75 484 transcript predictions. Most of the predictions showed a good overlap with the annotation (*F*_1_ ≥ 0.8). Nevertheless, 27 204 out of these 75 484 predictions had little (F_1_ <0.8) or no overlap with high or low confidence gene annotations. However, thousands of the transcripts contained in the official annotation do not have start or stop codons (Mascher *et al.*, 2017), which renders an exact comparison of predictions with perfect or at least very good overlap unreasonable.

Hence, we focus on 19 619 predictions with no overlap with any annotated transcript (Tab. 2). Scrutinizing these predictions, we find 1 729 single-coding-exon predictions that are completely covered by RNA-seq reads (tpc=1) but that are not contained in the annotation. Out of these, 367 are partially supported by best BLAT matches of ESTs to the genome. In addition, we analyzed multi-coding-exon predictions and find 2 821 predictions that obtain tie=1, stating that each predicted intron is supported by at least one split read from mapped RNA-seq data. Out of these, 1 390 are partially supported by best BLAT matches of ESTs to the genome.

Besides predictions that are well supported by RNA-seq data, we also observe thousands of predictions that are not (tpc = 0 or tie = 0) or only partially (0 < tpc < 1 or 0 < tie < 1) supported by RNA-seq. Despite no or only partial RNA-seq support, we find that 833 are partially supported by best BLAT matches of ESTs to the genome.

Alternatively, we can utilize the number of reference organisms that support a prediction (#evidence) to filter the predictions as noted for *D. melanogaster*. This approach will decrease sensitivity, but increase specificity obtaining predictions with a high confidence. Although, we find the most predictions with #evidence = 1, we also find about 3 500 predictions with #evidence *>* 1, more than 1 100 of these predictions are additionally supported by RNA-seq data or ESTs.

## 4 Conclusions

Summarizing the methods and results, we present an extension of GeMoMa that allows for the incorporation of RNA-seq data into homology-based gene prediction utilizing intron position conservation. Comparing the performance of GeMoMa with and without RNA-seq evidence, we demonstrate for all four organism included in the benchmark that RNA-seq evidence improves the performance of GeMoMa. GeMoMa performs equally well or better than BRAKER1, MAKER2, CodingQuarry, and purely RNA-seq-based pipelines on the benchmark data sets including plants, animals and fungi. In addition, we demonstrate that GeMoMa helps to improve the performance of combined gene predictor pipelines as for instance MAKER2. Notably, model organisms have been used as target organisms in this benchmark, whereas they would typically be used as reference organisms in real applications. Hence, the performance of homology-based gene prediction programs might be underestimated. In summary, we recommend to use homology-based gene prediction using RNA-seq data as implemented in GeMoMa whenever high-quality gene annotations of related species are available.

Interestingly, we find that GeMoMa works especially well for *D. melanogaster* in the benchmark study compared to the performance of its competitors. One possible reason could be that Flybase used homology and RNA-seq data besides other evidence to infer the gene annotation (Matthews *et al.*, 2015). In contrast, we find the worst results in *C. elegans* in the benchmark study, which might be related to the fact that the *C. elegans* gene set contains many rare isoform community submissions whereas *C. briggsae* was annotated by a large scale gene predictions effort based on RNA-seq.

Scrutinizing the annotation in Wormbase, we predicted protein-coding transcripts for four nematode species based on the annotation of the model organism *C. elegans*. We find that a substantial part of the GeMoMa predictions is either missing, marked as modification of annotated transcripts or alternative transcripts. Especially for the three nematodes, *C. brenneri*, *C. japonica* and *C. remanei*, that are annotated solely using ab-initio gene prediction, we find a large part of the annotation that is marked as questionable or missing. This may give an indication, why homology-based gene prediction for *C. elegans* shows less good performance in the benchmark study. The GeMoMa predictions of the four nematodes will be included in Wormbase in the upcoming releases. Furthermore, GeMoMa will be included in the WormBase gene curation process and trialled for strain annotation.

Furthermore, we predicted protein-coding transcripts for barley using four reference species and find several hundreds of predictions that are not included in the reference annotation but are supported by RNA-seq data, ESTs or multiple reference species. Hence, we conclude that these are valuable predictions harboring additional barley genes. These predictions will be incorporated in the new barley annotation.

GeMoMa provides a user-friendly documentation and can be use as command line tool or through the Galaxy workflow management system (Afgan *et al.*, 2016) providing its own Galaxy integration (Fig. S1). GeMoMa is freely available under GNU GPL3 at http://www.jstacs.de/index.php/GeMoMa.

## Acknowledgements

We thank Katharina Hoff for providing the BRAKER1 benchmark data sets, Carson Holt for assisting the MAKER2 comparison, Gil dos Santos for his comments on the quality of the Drosophila genome assemblies, Erich Schwartz for his comments on *C. japonica*, Gary Williams for his comments on *C. briggsae*, and Thomas Berner for technical assistance.

